# Single-cell metabolic oscillations are pervasive and may alleviate a proteome constraint

**DOI:** 10.1101/2024.11.25.625147

**Authors:** Arin Wongprommoon, Alán F. Muñoz, Diego A. Oyarzún, Peter S. Swain

## Abstract

Biological rhythms not only coordinate cellular activity with external signals, but may also enable internal coordination. The metabolic cycle in budding yeast is perhaps the most well-studied example. Historically researchers have investigated this cycle in populations growing in chemostats, but more recently time-lapse microscopy has revealed single-cell oscillations in the redox state of enzyme cofactors and in ATP levels. How to relate the results of these two types of assays is however unclear. Here we report single-cell rhythms too in intracellular pH and show that oscillations in the redox state of flavin molecules occur in auxotrophic and prototrophic strains, in nutrients favouring respiration or fermentation, and in deletion mutants for which oscillations in chemostats are either unobservable or disrupted. To explain the pervasiveness of these rhythms, we postulate that cells generate oscillations to alleviate a proteome constraint – amino acids cells use for one class of enzymes are unavailable for others. Using flux balance analysis with an enzyme-constrained genome-scale metabolic model, we show that, with a finite proteome, sequential synthesis of biomass components typically generates a shorter doubling time than synthesising components in parallel. Our results suggest that the metabolic cycle drives growth and is potentially widespread because all cells grow within a proteome constraint.

## Introduction

Biological rhythms are widespread and (Glass, 2001) are typically associated with an external signal. For example, circadian rhythms coordinate organisms’ activities with the daily light-dark cycle. As well as external coordination, it has been reported that biological rhythms might also enable internal coordination and allow intracellular events to follow a fixed temporal sequence (Millar, 2016).

The single-celled eukaryote budding yeast provides perhaps the best studied example of internal coordination. If grown at high density in aerobic, nutrient-limited chemostats, populations of budding yeast cells can spontaneously synchronise and undergo metabolic oscillations (Mellor, 2016), generating periodic rhythms in transcription (Klevecz et al., 2004; Tu et al., 2005) and in levels of metabolites (Murray et al., 2007; Tu et al., 2007a). A defining characteristic of metabolic oscillations is periodic changes in dissolved oxygen in the medium (Mellor, 2016).

Since metabolic oscillations could be artefacts of synchronised cell populations, early works studied rhythms in single-cell transcriptional patterns (Silverman et al., 2010). With the adoption of microfluidic technology and long-term imaging, several works reported single-cell oscillations in key metabolic readouts, such as the redox state of nicotinamide nucleotide NAD(P)H (Papagiannakis et al., 2017) and flavin (Baumgartner et al., 2018), as well as ATP levels (Papagiannakis et al., 2017), typically one-to-one with the cell cycle.

Nevertheless how these single-cell rhythms relate to those observed in chemostats and why metabolism should oscillate at all are both unclear. Preventing respiration does not stop single cells from oscillating (Baumgartner et al., 2018) despite changing oxygen consumption being considered an essential feature of population-level studies. Cells might generate their rhythms through amassing stores of carbohydrates (O’ Neill et al., 2020) and then liquidating these stores at once to power through the cell cycle (Futcher, 2006). Yet single-cell rhythms persist in mutants unable to generate both glycogen and trehalose (Takhaveev et al., 2023).

Here we use microfluidics and time-lapse microscopy to monitor cycles of flavin redox-state in budding yeast. We show that single-cell rhythms persist in auxotrophic as well as in prototrophic strains, in different nutrient conditions, and in deletion mutants whose oscillations in chemostats are either unobservable or perturbed. The oscillations are usually one-to-one with the cell cycle, which we follow simultaneously using a fluorescently tagged histone. Given this prevalence, we conjecture that the rhythms are fundamental to growth and arise as a mechanism to alleviate a proteome constraint. Using flux balance analysis with an enzyme-constrained genome-scale metabolic model, we test this hypothesis *in silico* and demonstrate that with a finite proteome synthesising biomass components sequentially typically generates shorter doubling times than synthesising in parallel. If this sequence has a fixed order, it will manifest as a metabolic rhythm. Our work provides new insights on the origins of metabolic oscillations in single-cells and suggests that they may be a widespread strategy to mitigate proteome constraints.

## Results

### Cells have flavin-redox oscillations typically phase-locked with the cell cycle

Following Baumgartner et al. (2018), we monitored the redox state of flavin molecules — riboflavin, flavin mononucleotide (FMN), and flavin adenine dinucleotide (FAD) — by measuring their collective autofluorescence in individual cells. These molecules fluoresce in the visible spectrum, more when oxidised and less when reduced (Chance & Schoener, 1966). Baumgartner et al. (2018) reported flavin-redox oscillations as typical of single cells. For our experiments, we performed time-lapse fluorescence microscopy using ALCATRAS microfluidic devices (Crane et al., 2014) and the BABY algorithm to automatically segment, track cells over time, and predict budding events from the bright-field images (Pietsch et al., 2023).

We postulated that if cycling cells of budding yeast simultaneously undergo metabolic cycles, then we should observe metabolic cycles in auxotrophic strains as well as in the prototrophic strains studied previously (Baumgartner et al., 2018; Papagiannakis et al., 2017). Using our setup, we measured robust flavin-redox oscillations in cells of the BY4741 strain (Fig. 1A), a standard, auxotrophic laboratory strain used for both the budding yeast deletion and GFP collections (Giaever et al., 2002; Huh et al., 2003).

**Figure 1.**
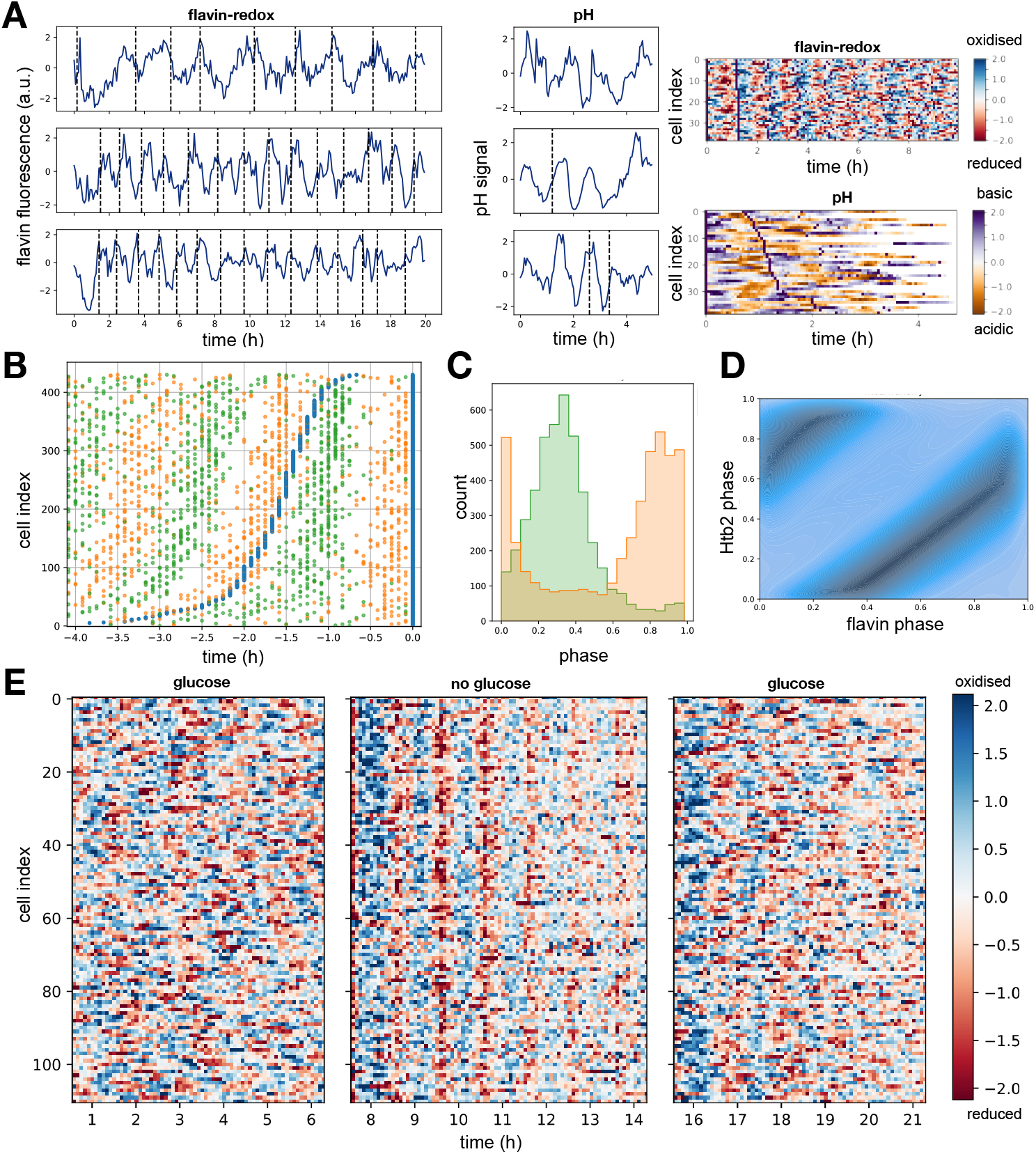
Single cells have oscillations in the redox state of flavin molecules that typically phase-lock with the cell cycle. **A** Single-cell rhythms in the flavin-redox state and intracellular pH indicate underlying oscillations in metabolism, even in BY4741 auxotrophs. BY4741 cells were in 1% glucose, while pHluorin cells were in 2% glucose. Dotted lines mark budding events. For the heat maps, we aligned cells by their first budding event to highlight the oscillations. Navy dots mark subsequent buddings. We normalised the fluorescence data for each cell to have a mean of zero and measured pH using the pH-sensitive Green Fluorescent Protein pHluorin2 (Mahon, 2011). **B** A raster plot shows the sequence of cell-cycle events between two consecutive peaks in the flavin-redox oscillations, here for the prototrophic FY4 strain. We aligned cells using their flavin cycle closest to *t* = 8h into the experiment, a time arbitrarily chosen. Blue marks peaks in the chosen flavin oscillation – we do not show earlier flavin oscillations for clarity; orange marks budding events, the start of S phase; green marks peaks in Htb2-mCherry fluorescence, M phase. **C** Defining the phase of each signal to be zero at one peak and to approach one at the subsequent peak, both budding events (orange) and maximal Htb2 fluorescence (green) occurred at distinct points in the flavin cycle. We binned data for all cells. **D** A kernel-density plot of the single-cell flavin and cell-cycle phases using data from all cells shows phase-locked behaviour, with the dark, highly populated regions indicating a stable attractor. **E** A heatmap of flavin-redox states across a population of cells indicates that the flavin cycles, although asynchronous during high-glucose conditions (0.75% in minimal medium), became synchronous during six hours of glucose starvation when most cells slowed or arrested their cell cycle.

Consistent with a metabolic cycle generating these rhythms we also observed single-cell oscillations in intracellular pH using a pH-sensitive version of GFP Mahon, 2011 (Fig. 1A). These oscillations had a period typically equal to the cell cycle (Fig. S3). Population-level oscillations in intracellular pH occur too in metabolically synchronised cells growing in continuous culture (O’ Neill et al., 2020). In exponentially growing yeast, the plasma membrane protein Pma1 consumes ATP to keep the intracellular pH near neutral by exporting protons (Kane, 2016). ATP levels are known to oscillate one-to-one with the cell cycle (Papagiannakis et al., 2017), and the oscillations of intracellular pH we observed might reflect periodic changes in ATP availability altering Pma1’s activity. Alternatively, they might be indirectly driven by changes in vacuolar pH, which can oscillate with the cell cycle because cells regulate the storage and release amino acids through vacuolar pH (Okreglak et al., 2023).

To confirm the reported phase relationship between the flavin-redox oscillations and the cell cycle (Baumgartner et al., 2018), we used mCherry to tag a histone gene, HTB2, in the FY4 prototrophic strain. The Htb2 signal peaks in M phase (Garmendia-Torres et al., 2018) as the cell replicates its genome. Combining these measurements with the budding events identified by the BABY algorithm, which mark the start of S phase (Hartwell et al., 1974), we found that both S and M phase occur at distinct phases of the flavin-redox cycle (Fig. 1B). The data also reveals that peaks in flavin signal, which we define arbitrarily to mark the beginning and end of its cycle, occur at budding events, while the M phase, where the Htb2 signal is maximal, occurs approximately at the end of the flavin cycle’s first quarter (Fig. 1C). The flavin signal is likely minimal early in G1 (Fig. S1). These observations imply that the two cycles are one-to-one phase-locked. Plotting the cell-cycle phase against the flavin-redox phase gives the classic phase portrait of two coupled, phase-locked oscillators (Strogatz, 1994) (Fig. 1D). Similar coupling has been reported between the cell and the circadian cycles in mammalian cells (Feillet et al., 2014).

We observed similar behaviours, although with a longer G1 phase and larger amplitudes, when we changed central carbon metabolism by growing cells in pyruvate (Fig. S1), which, unlike glucose, cells cannot metabolise via fermentation but only via respiration (Fraenkel, 2011). Chemostat experiments are often performed with low dilution rates (Mellor, 2016), generating, for example, metabolic cycles with long periods of five to eight hours (O’ Neill et al., 2020), and so we also tried experiments with minimal (0.001%) glucose. The oscillations however, if present, had such low amplitudes and such low signal-to-noise ratios that they were too difficult to detect reliably (Fig. S5) even though the cell-cycle times were similar to those in 2% pyruvate.

Previous work has shown that although cycling cells without metabolic oscillations are typically not observed, arrested cells with metabolic oscillations are (Baumgartner et al., 2018; Özsezen et al., 2019; Papagiannakis et al., 2017). To study the interplay between the two cycles, we decoupled them by abruptly switching media to deprive cells of glucose. Most cells then arrested or slowed their cell cycle (Fig. S2A & B), but not only did the flavin-redox oscillations continue, they also synchronised across the cells (Fig. 1E), with their amplitude falling over time. Similarly, intracellular pH oscillations also continued during glucose starvation in a separate experiment (Fig. S2C). When we switched cells back into glucose (Fig. 1E), normal amplitude oscillations resumed, and synchrony was gradually lost. These observations show that metabolic oscillations can reset their phase in response to nutrient changes and support the proposal that they are autonomous and not necessarily coupled to the cell cycle (Baumgartner et al., 2018; Papagiannakis et al., 2017).

Taken together, our results confirm an earlier report of robust flavin-redox oscillations measured in single yeast cells with a different genetic background (Baumgartner et al., 2018). The oscillations are typically phase-locked one-to-one with the cell cycle (Fig. S3), but can persist in non-cycling cells. We observed oscillations too in intracellular pH, suggesting that both these rhythms are features of an underlying oscillation that pervades metabolism (Baumgartner et al., 2018; Papagiannakis et al., 2017).

### Flavin-redox oscillations are more robust to genetic perturbations than population-level rhythms

Early studies focused on metabolic oscillations in populations of yeast cells, synchronising their behaviour by an initial period of starvation and then monitoring their changing and periodic consumption of oxygen (Mellor, 2016). A key finding was that specific gene deletions can cause the oscillations to become undetectable or alter their period or phase (Causton et al., 2015).

To investigate whether the flavin-redox oscillations are the same metabolic rhythms studied in continuous culture although manifested in single cells, we studied two strains reported to have perturbed oscillations in chemostats: the *zwf1*Δ strain, which does not generate oscillations in dissolved oxygen unlike its parental strain (Tu et al., 2007b), and the *tsa1*Δ *tsa2*Δ strain, which does oscillate but with a waveform having two peaks per cycle rather than the usual one (Amponsah et al., 2021; Causton et al., 2015).

For both these strains, we observed oscillations in the redox state of flavin molecules, and they remained one-to-one phase-locked with the cell cycle (Fig. 2 & S3). Although the cell-cycle period and the flavin-redox cycle period increased in the *zwf1*Δ strain relative to the wild-type (Fig. 2A & S3), it robustly oscillated with a waveform indistinguishable from the wild-type (Fig. 2B), in contrast to the reported behaviour in chemostats (Tu et al., 2007b). Compared to the wild type, the *tsa1*Δ *tsa2*Δ strain had oscillations with a weaker amplitude, more variable period (Fig. 2C & S3), and a perturbed phase (Fig. 2D), but they did not have any characteristics analogous to the two peaks per cycle seen in chemostats.

**Figure 2.**
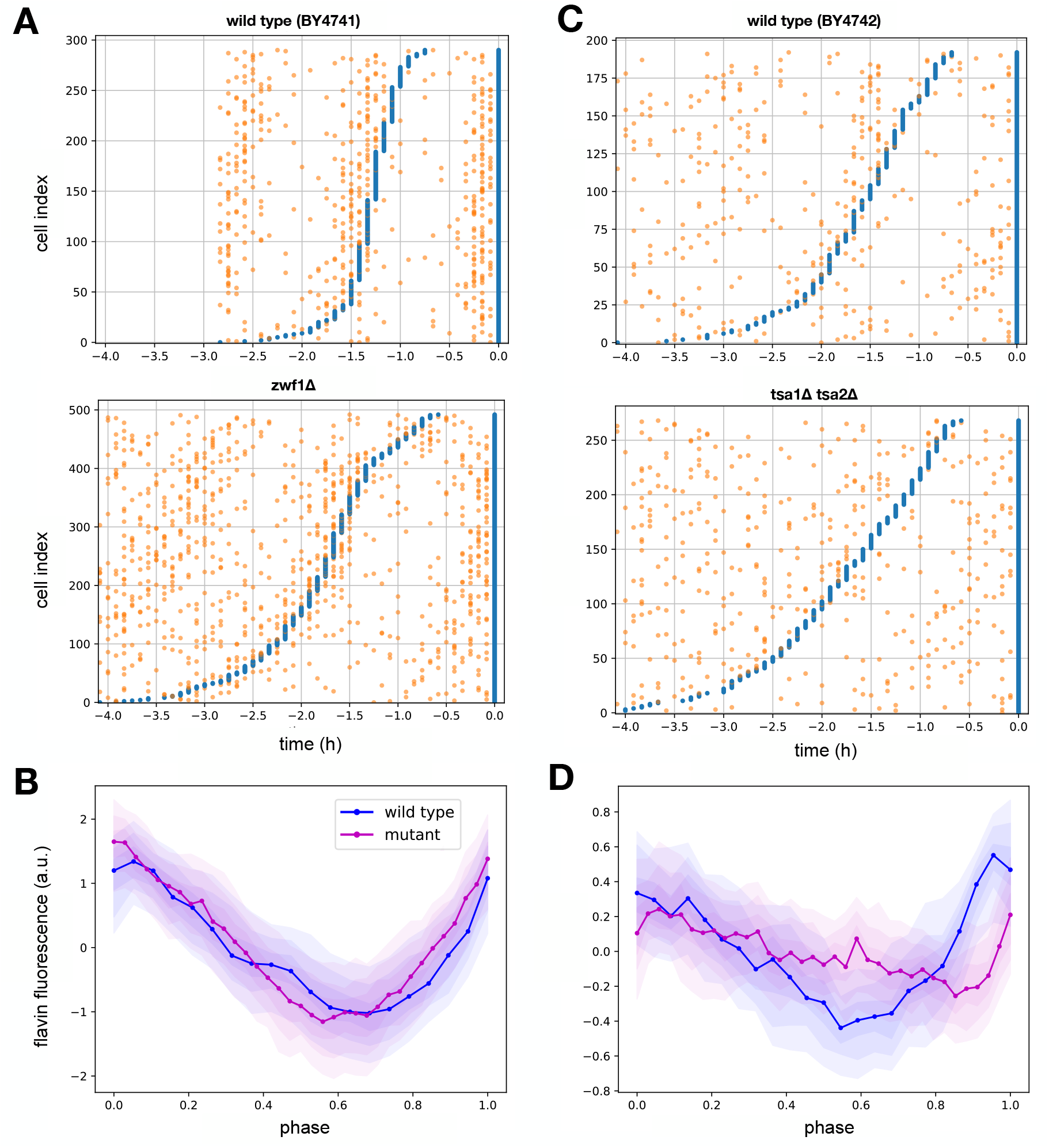
Gene deletions that perturb metabolic oscillations in chemostats only weakly affected single-cell flavin-redox rhythms. **A** Raster plots comparing the flavin-redox oscillations between the *zwf1*Δ and its BY4741 parent strain. Oscillations persisted and had a longer period. **B** Averaging the single-cell flavin cycles between two consecutive peaks showed that the waveform of the *zwf1*Δ mutant (purple) is indistinguishable from that of its parent BY4741 strain (blue). We aligned the data at *t* = 9h. **C** Raster plots comparing the flavin-redox oscillations between the *tsa1*Δ *tsa2*Δ strain and its BY4742 parent strain. Oscillations were noisier in the mutant, with the second region mostly free of budding events in the parent strain, around *t* = *−*2.2h, obscured in the mutant. **D** The waveform of the *tsa1*Δ *tsa2*Δ strain (purple) has both a smaller amplitude and dips later in the cycle than the waveform of its parent strain (blue). We aligned the data at *t* = 9h.

As a further test, we replaced potassium in 2% glucose medium with sodium, a change that when implemented gradually in chemostats decreased the amplitude of the oscillations in dissolved oxygen while increasing their period until the oscillations disappeared (O’ Neill et al., 2020). In our experiments, we could only make abrupt extracellular changes and such a change in potassium availability had little effect (Fig. S4). We saw robust oscillations, although with more variability in the phase of the flavin cycle at the Htb2-fluorescence peaks.

### Alleviating a proteome constraint by sequentially synthesising biomass can generate metabolic rhythms

The persistence of rhythms in the flavin-redox state across nutrient and genetic perturbations suggests that the metabolic cycle is an intrinsic property of a growing yeast cell. To investigate why, we sought to explain the oscillations as a consequence of cells temporally ordering their biosynthesis to alleviate constraints on intracellular resources.

Microbes have limited resources (Scott & Hwa, 2022), including ATP, biosynthetic machinery, physical space (Elsemman et al., 2022), and particularly amino acids (Scott et al., 2010). Dedicating amino acids to one class of proteins reduces amino acids available for other classes. To grow and reproduce, cells should appropriately allocate amino acids between anabolic and catabolic enzymes (Scott & Hwa, 2022; You et al., 2013), preventing excessive catabolism limiting anabolism and vice versa.

With limited numbers of enzymes, we postulated that a cell might more efficiently generate the constituents of new cell by dedicating its entire pool of free amino acids to synthesise each class of constituents one by one, rather than simultaneously synthesising all together. The re-routing of amino acids, say between the enzymes for synthesising carbohydrates to those for synthesising lipids, will change metabolic activity. If cells synthesise the different constituents in a fixed order, this changing metabolic activity will manifest as periodic metabolic oscillations that are one-to-one with the cell cycle and constitutive to growing cells (Fig. 3A).

**Figure 3.**
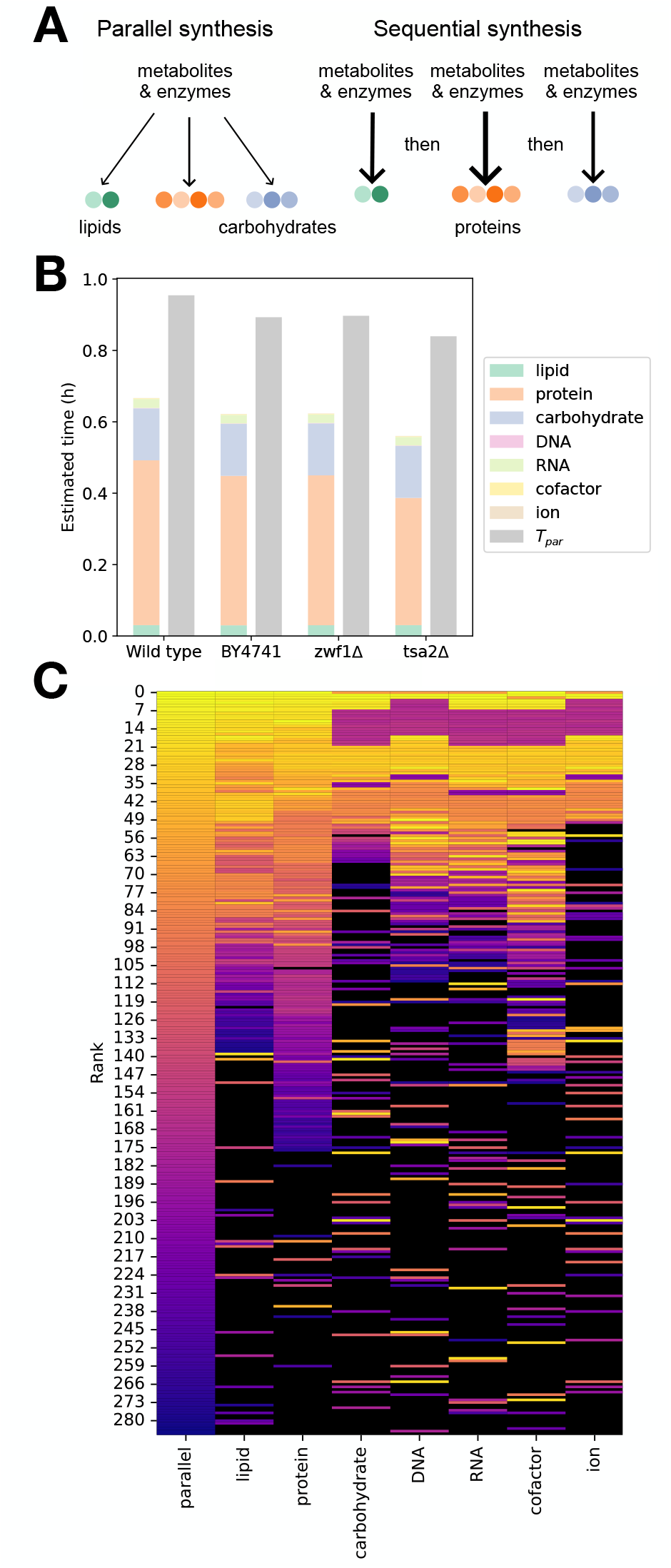
A limited proteome pool can favour sequential, rather than parallel, synthesis of the components of a new cell, with the re-partitioning of amino acids from one synthesis process to the next potentially generating metabolic rhythms in growing cells. **A** For parallel synthesis, cells arrange their proteome to synthesise all biomass components simultaneously; for sequential synthesis, cells dedicate the proteome to synthesising each class of biomass constituents in turn. **B** Using flux balance analysis and the ecYeast8 model (Lu et al., 2019), we compared the total time to synthesise seven biomass components sequentially (left bars) to the time to synthesise all in parallel (right bars). In the rich medium considered here, with glucose imported at a rate of 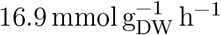, sequential synthesis generated shorter doubling times and so faster growth. **C** Sequential synthesis is likely faster because cells allocate their proteome differently when synthesising only carbohydrates to when they synthesise only proteins, for which their proteome allocation is similar to parallel synthesis. In each column, rows represent enzymes, ranked in descending order of their numbers (strictly, 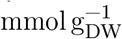). Colours identify the ranking for parallel synthesis, with black indicating missing enzymes 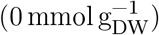. How much the order changes indicates the extent to which the proteome allocation differs from parallel synthesis.

To test this hypothesis, we employed genome-scale metabolic models to simulate and compare temporal strategies. Using the ecYeast8 enzyme-constrained metabolic model of budding yeast (Lu et al., 2019), we were able to constrain the number of metabolic enzymes and predict the maximal growth rate, as quantified by a biomass reaction. In the ecYeast8 model, biomass has seven components (Lu et al., 2019): lipids, which include sterols, fatty acids, triacylglycerols, and phospholipids; proteins, including all amino acids; carbohydrates, such as beta-D-glucans, mannan, glycogen, and trehalose; DNA, including nucleosides; RNA; cofactors, such as coenzyme A, FAD, NAD(P), thymidine diphosphate, and tetrahydrofolic acid; and ions, such as metal cations, sulphate, and chloride. These distinct components of biomass allowed us to predict growth rates for both sequential and parallel synthesis.

For sequential synthesis, we altered the biomass reaction so that it generated only one of its components, taking each in turn. To determine the time to synthesise sufficient component *i* for a new cell, *t*_seq,*i*_, we used the maximal flux of the biomass reaction when the cell synthesises only component *i, λ*_seq,*i*_, and the fraction of the biomass comprising *i, f*_*i*_, so that

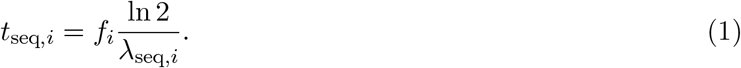

We defined the sequential doubling time as

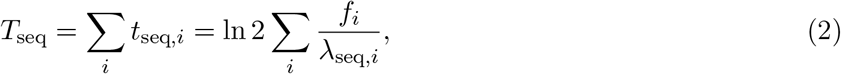

the sum of the times to synthesise each component. For example, if all the *λ*_seq,*i*_ are equal, *T*_seq_ becomes ln 2*/λ*_seq,*i*_ because the *f*_*i*_ sum to one by definition (Table S7).

For parallel synthesis, we used ecYeast8’s full (unmodified) biomass reaction and computed the synthesis time for the biomass components by assuming the time for each to be proportional to its mass fraction and inversely proportional to the maximal growth rate *λ*:

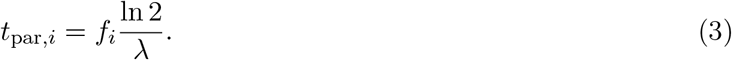

We then defined the parallel doubling time as the longest of these synthesis times:

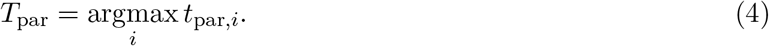

For ecYeast8, almost 90% of dry mass is protein and carbohydrate, with *f*_protein_ *> f*_carbohydrate_ (Lu et al., 2019) (Table S7), and so *T*_par_ = *t*_par,protein_.

Comparing the two strategies using the ratio of the sequential doubling time to the parallel one, *τ*seq*/*par = *T*_seq_*/T*_par_, we found sequential biosynthesis to be faster than parallel biosynthesis when cells rapidly uptake glucose (Fig. 3B). To understand why, we visualised the change in enzyme use through ranking the enzymes by the amount of the proteome they received during parallel biosynthesis and determining how these ranks changed for sequential synthesis. The proteome allocation when synthesising only proteins is similar to that when synthesising only lipids and both are similar to the proteome allocation during parallel synthesis (Fig. 3C). When synthesising only carbohydrates and the remaining biomass components, although the protein allocation is similar across components, it differs from parallel synthesis. Sequential synthesis is likely favoured overall because the proteome allocation for the synthesis of proteins and for the synthesis of carbohydrates, the two largest biomass components, show such markedly different patterns. This difference undermines parallel synthesis because the cell cannot synthesise both components with the same enzymes but instead must allocate enzymes to each, reducing the amount synthesising proteins, and so increasing *t*_par,protein_ and *T*_par_.

Furthermore, by deleting the appropriate enzymes from the model, we showed that sequential synthesis is fastest too for metabolic networks that mimic the strains in which we saw flavin-redox rhythms: the auxotrophic BY4741 strain and the *zwf1*Δ and *tsa2*Δ deletion strains — there is no TSA1 gene in ecYeast8 (Fig. 3B).

### A constrained proteome usually favours sequential biosynthesis

To explore when sequential synthesis is preferred, we simulated different availabilities of both nitrogen, by varying the ammonium exchange rate, and carbon, by varying the exchange rate for either glucose or pyruvate with each as the only carbon source. We found that parallel synthesis could become faster than sequential synthesis (Fig. 4A), particularly when both carbon and nitrogen limited growth and to a lesser extent when nitrogen weakly limited growth but carbon did not.

**Figure 4.**
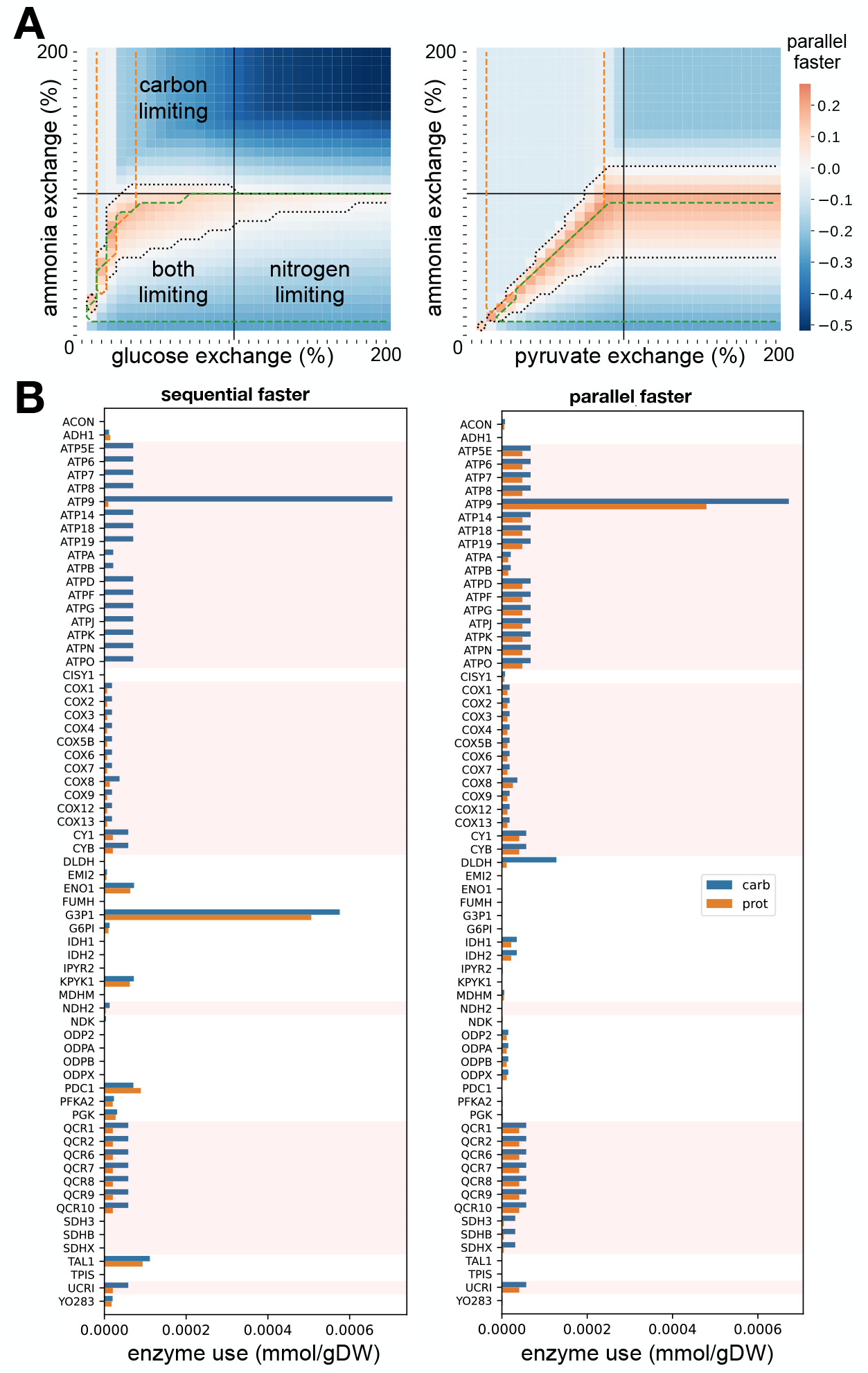
Sequential synthesis is usually but not always better than parallel synthesis. **A** A heatmap of log_2_(*τ*_seq*/*par_) for varying glucose and ammonium exchange rates showing where parallel synthesis is faster (red region). Exchange rates are given as percentages of the exchange rate that maximises the growth rate. To determine limiting regions, we calculated the logarithmic sensitivity of the maximal growth rate to either the carbon or ammonium exchange rate (Ingalls, 2008). If carbon sensitivity is small (*<* 0.01), then nitrogen is limiting; if nitrogen sensitivity is small (*<* 0.01), than carbon is limiting; if both sensitivities are small, neither are limiting; and if neither sensitivity is small, then both are limiting. **B** When sequential synthesis is faster, there are greater changes in enzyme use from when the cell synthesises only protein biomass to synthesising only carbohydrate biomass. We show the enzyme numbers for enzymes that participate in generating both carbohydrate and protein biomass. Those enzymes with background shading contribute to the electron transport chain and so respiration. Their use changes between generating carbohydrate and protein biomass if sequential synthesis is best.

The times to synthesise the protein and carbohydrate biomass components mainly determine the better strategy. For sequential synthesis to be faster, *T*_seq_ *< T*_par_ implies, from their definitions,

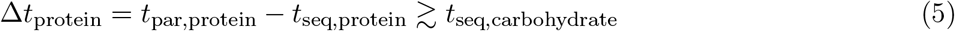

so that the reduction in time generating protein biomass through sequential synthesis must be greater than the time to generate carbohydrate biomass.

We expect sequential synthesis to be faster if carbon is plentiful. When synthesising protein biomass, the cell still requires central carbon metabolism to provide energy, reducing power, and carbon backbones for amino acids (Fraenkel, 2011). If carbon is abundant, its availability will saturate some enzymes and make their flux greater than required. Consequently the cell can move resources from these enzymes towards other biomass-generating processes while still maintaining sufficient carbon flux, thus favouring sequential synthesis. Such a move could be through changes in gene expression or in rates of enzyme degradation. In contrast, when both carbon and nitrogen are limiting, fewer if any fluxes will be in excess. The cell has little freedom to reallocate resources: doing so for one of the required processes will decrease that process’s flux even if increasing another’s. Sequential synthesis therefore becomes disfavoured.

To observe this phenomenon, we focused on enzymes that generate both protein and carbohydrate biomass: for these enzymes we can unambiguously compare how their levels change because they never become zero. When the extracellular conditions favoured sequential synthesis, we typically observed a substantial change in enzyme use when synthesising proteins compared to carbohydrates (Fig. 4B). The extracellular conditions alleviated trade-offs in reallocating proteome among these shared enzymes, allowing the cell to grow faster. In conditions favouring parallel synthesis, however, the enzymes typically had similar levels when synthesising proteins and carbohydrates (Fig. 4B), implying these conditions hinder reallocation.

We found that when conditions favour sequential synthesis, the levels of enzymes in the electron transport chain differs between protein and carbohydrate synthesis. This difference suggests that, in our model, cells favour respiration when generating carbohydrate biomass and fermentation when generating protein biomass. Even with rapid glucose import (Fig. 3A), we observed that cells synthesising carbohydrate biomass consume more than twice as much oxygen and produce 30% less carbon dioxide and 20% less ethanol compared to cells synthesising protein biomass. This changing importance of respiration versus fermentation as cells synthesise these two biomass components, with the accompanying changes in oxygen consumption, is strongly suggestive of the changing, periodic rates of oxygen consumption used to identify metabolic oscillations in chemostats.

## Discussion

Building on earlier work (Baumgartner et al., 2018; Papagiannakis et al., 2017), we have shown that auxotrophic and prototrophic single cells have robust rhythms in their flavin-redox state in environments favouring either fermentation or respiration.

These rhythms persist in deletion strains that have disrupted metabolic oscillations when assayed as a population, with their rate of oxygen consumption in chemostats distinct from the wild-type strain’s (Amponsah et al., 2021; Causton et al., 2015; Tu et al., 2007b). To be able to measure rhythms in populations, cells not only must oscillate but also oscillate in synchrony, and metabolically oscillating cells in continuous culture likely have not one but multiple synchronous subpopulations (Slavov et al., 2011). Although our results for the *zwf1* Δ and *tsa1* Δ *tsa2* Δ strains differ from the equivalent chemostat experiments, the two can be reconciled if the deletions do not principally disrupt the rhythms but the cell-to-cell synchrony.

Our work suggests that metabolic oscillations exist in growing cells and are typically phase-locked one-to-one with the cell cycle in cells that both grow and divide. A caveat is that the single-cell measurements are noisy, and at times it was challenging to determine whether or not a rhythm existed. For example, we were unable to unambiguously identify flavin-redox oscillations for cells growing in minimal medium with 0.001% glucose. We suspect that this failure is likely because of the data’s low signal-to-noise ratio rather than an absence of rhythms: the amplitude of the signal we do find is about half that in 2% glucose.

The design of our microfluidic device, with cells growing in flowing medium in their own distinct, physically separated “jails” (Crane et al., 2014), makes interactions between cells via the medium unlikely, yet we still saw metabolic rhythms. This observation casts doubt on previous suggestions that oscillations may result from cell-to-cell communication (Mellor, 2016).

Instead, to explain the prevalence of metabolic rhythms in single cells, we postulated that a limited proteome might make it more effective for cells to synthesise the different constituents of a new cell sequentially rather than simultaneously. In contrast to having co-existing but lower levels of all the necessary enzymes, faster synthesis might result if the cell reallocates amino acids from one class of enzymes to another as it generates the different components of biomass. If there is a fixed sequence of synthesis, then metabolism will shift periodically from one type of synthesis to the next as cells grow and divide, generating a rhythm phase-locked to the cell cycle (Takhaveev et al., 2023). Cells do appear to have a temporal order as they synthesise biomass, generating protein mainly in G1 (Campbell et al., 2020; Guerra et al., 2022; Litsios et al., 2019; Takhaveev et al., 2023) and lipids in S and G2/M phases (Blank et al., 2020; Campbell et al., 2020; Takhaveev et al., 2023). Using enzyme-constrained flux balance analysis, we demonstrated that fixing the size of the proteome can make sequential synthesis faster than parallel synthesis and usually does so for the conditions we investigated. The same approach predicts oscillations in dissolved oxygen for synchronised populations because the cells favour fermentation when generating protein biomass and respiration when generating carbohydrates.

One prediction is that cells that undergo multiple flavin oscillations in one cell cycle should increase in volume more than cells that undergo a single flavin oscillation. Re-analysing the data from Fig. 1E, we did see a weak positive correlation between a cell’s change in volume and the number of flavin cycles per Htb2 cycle while cells entered and recovered from starvation (Fig. S6). These cells were however in minimal medium, and so any growth likely came from stored carbohydrates. Growth does continue robustly with NAD(P)-redox oscillations for cells genetically arrested in glucose (Papagiannakis et al., 2017). Our data suggest that the changes in flavin-redox state indicate that cells are repeatedly poised to synthesise biomass throughout the starvation period, perhaps using nutrient-sensing networks to determine whether they should launch a wave of growth.

Taken together, our findings support earlier speculations that the metabolic cycle predominately controls growth and that the cell cycle controls division (Slavov et al., 2011). We have shown that metabolic rhythms can arise through cells alleviating a proteome constraint, an example of a universal restriction on growth. Elucidating the logic of the yeast metabolic cycle may therefore reveal the origin of biological rhythms more broadly.

## Materials and Methods

### Strains and growth media

The *Saccharomyces cerevisiae* strains we used are in Table 1.

**Table 1.** Strains used in this study.

| Name | Background | Genotype | Origin | Notes |
| --- | --- | --- | --- | --- |
| FY4 | FY4 | — | EUROSCARF | Winston et al. (1995) |
| htb2::mCherry | FY4 | HTB2::mCherry | In-house, CRISPR | — |
| BY4741 | BY4741 | *MATa his3Δ1 leu2Δ0 met15Δ0 ura3Δ0* | EUROSCARF | Brachmann et al. (1998) |
| *zwf1Δ* | BY4741 | *zwf1Δ::KAN* | Edinburgh Genome Foundry | Yeast deletion collection |
| BY4742 | BY4742 | *MATα his3Δ1 leu2Δ0 lys2Δ0 ura3Δ0* | Bruce Morgan | Calabrese et al. (2019) |
| pHluorin | BY4742 | pAG415GPD-pHluorin2 | Agnes Toth-Petroczy | Munder et al. (2016) |
| *tsa1Δ tsa2Δ* | BY4742 | *tsa1Δ::natNT2 tsa2Δ::kanMX4* | Bruce Morgan | Calabrese et al. (2019) |
| CEN.PK113-7D | CEN.PK113-7D | — | Peter Kötter | Nijkamp et al. (2012) |

We used the minimal medium described by Verduyn et al. (1992) unless otherwise stated, adjusting its pH to 6.0 using potassium hydroxide, or sodium hydroxide for potassium-free media. The exception was the pHluorin strain, for which we used synthetic complete medium.

For BY4741-background strains, the growth medium was supplemented with 125 mg L^−1^ histidine, 500 mg L^−1^ leucine, 75 mg L^−1^ tryptophan, 100 mg L^−1^ methionine, and 150 mg L^−1^ uracil (Pronk, 2002). For BY4742-background strains, the growth medium was supplemented similarly, but replacing methionine with 100 mg L^−1^ lysine-HCl. We added too a carbon source.

### Single-cell microfluidics

To prepare cell cultures, we grew yeast strains from glycerol stocks on solid agar. Then, single colonies were added to liquid culture in minimal media, with supplements for auxotrophic strains and a carbon source. For the high glucose, potassium-deficiency, and pH-oscillation experiments, we used 2% (20 g L^−1^) glucose. For the glucose-starvation experiment, we used 0.75% glucose; for the low-glucose experiment, 0.001% glucose; for pyruvate experiments, 2% sodium pyruvate; and for experiments with deletion strains, 1% glucose. We incubated liquid cultures at 30 °C for either 14 h if the carbon source was glucose or 48 h if the carbon source was pyruvate. The cells were then diluted again so that the resulting culture had an OD_600_ of 0.10–0.20 and incubated for a further 4 h.

To monitor metabolic cycles in single cells, we used ALCATRAS microfluidics (Crane et al., 2014). To prepare for an experiment, a device’s chambers were filled with growth medium supplemented with 0.05% w/v bovine serum albumin and cells loaded. To control nutrient conditions, we connected the device to two syringes containing different media and programmed the ALCATRAS system to switch between the syringes for the glucose starvation, potassium-deficiency, and pH oscillation experiments. The resulting flow rate was approximately 4 µL min^−1^. The cells, ALCATRAS chambers, and syringes were kept in a 30 °C incubation chamber (Oko-labs).

Microscopy was performed using a 40 *×* 1.4 NA oil immersion objective (Nikon), and the Nikon Perfect Focus System was used to ensure consistent focus. X-Y spatial positions were defined for each chamber to maximise spatial coverage of the chamber while ensuring that the microscope took less time to move positions and capture images than the interval period. Images were taken every 5 min for all experiments, except for the pH oscillation experiment in which images were taken every 3 min. For all strains, brightfield images were captured. Fluorescence imaging was performed with an OptoLED light source (Cairn Research), and LED voltage was optimised for maximum signal intensity without LED cut-off prior to experiments. Flavin (excitation 430/24, emission 535/30, exposure 60 ms) images were captured for all experiments except for the pH oscillation experiment, in which GFP (excitation 405/20, emission 480/40, exposure 30 ms) images were captured instead. In addition, mCherry (excitation 572/35, emission 632/60, exposure 100 ms) images were captured for the HTB2::mCherry strain. Five z-slices were taken for brightfield, GFP, and mCherry images, with a spacing of 0.6 µm between slices.

### Image analysis

To process the microscopy images, we used the `BABY` (Pietsch et al., 2023) algorithm within the `aliby` software package (Muñoz González, 2023), an end-to-end Python-based pipeline for time-lapse microscopy. With `BABY` to segment, track cells, and estimate single-cell growth rates, `aliby` produced time series for all cells of growth rate, volume, and background-corrected fluorescence for each fluorescence channel.

### Time-series analysis

To remove slow fluctuations that may confound our analysis, we applied a high-pass Butterworth filter with a critical frequency corresponding to a period of 350 min to all fluorescence time series.

We used `scipy`’s `find_peaks`, with some embellishments (given below), to identify peaks in each time series and so periods as peak-to-peak times. From `BABY`’s predicted budding events — a binary signal for each cell, we first estimated the median bud-to-bud time to approximate an experiment’s typical cell-cycle duration. We used this bud-to-bud time to process predicted peaks.

To estimate peaks in the flavin-redox signal, we applied a low-pass filter with a cut-off of 25% of an experiment’s Nyquist frequency – we typically sample every five minutes, but sampled every three minutes for the pHluorin experiment. We then used `find_peaks` with a `prominence` of 2.5 and a `distance` of the larger of 20 minutes or the bud-to-bud time divided by 12, both parameters being chosen by eye.

For the Htb2 signal, we developed an approach that exploited the signal’s saw-tooth pattern. First we normalised so that the signal’s minimum was zero and maximum one and ran `find_peaks` with a prominence of 0.01. The Htb2 signal falls sharply at cell divisions. For each peak, we therefore calculated its right prominence, the prominence for future times only, and the size of the drop from the peak to the minimal value of the signal before the next peak. We chose peaks with a right prominence above 0.3 and a drop above 0.3. Using these peaks, we created a binary signal and ran `find_peaks` again on this signal with a `distance` of a quarter of the bud-to-bud time to eliminate false positives.

For the pHluorin signal, we normalised so that the signal’s minimum was zero and maximum one and ran `find_peaks` with a prominence of 0.25 and a `distance` of the larger of 20 minutes or the bud-to-bud time divided by four.

Finally we ran `find_peaks` on BABY’s predicted budding events with a `distance` of a third of the bud-to-bud time to remove false positives.

We note that noise in the data affected the automated estimation of the peaks and periods of the signals, which we found as peak-to-peak times. Erroneously identified and missed peaks inevitably influenced our analysis and contributed to the spread of the distributions in Fig. S3.

### Flux balance analysis

For flux balance analysis, we used the ecYeast8.6.0 model (Lu et al., 2019) and COBRA (Python).

The model was modified to simulate non-wild type strains. The auxotrophy of BY4741 was simulated by allowing uptake of histidine, leucine, tryptophan, methionine and uracil, while BY4742 was simulated in a similar way but with lysine uptake replacing methionine uptake. Deletion of *ZWF1* and *TSA2* was performed using the gene-protein map in the ecYeast8 model; *TSA1* deletion was not performed for the *tsa1*Δ *tsa2*Δ strain because the model did not include reactions that correspond to this gene.

To simulate producing each class of biomass component in turn, we ablated pseudometabolites from the biomass reaction. To explain the process, consider the objective function, the biomass reaction:

~~~
     47.6 atp_c + 47.6 h2o_c + lipid_c + protein_c + carbohydrate_c
+ dna_c + rna_c + cofactor + ion
–> 47.6 adp_c + biomass_c + 47.6 h_c + 47.6 pi_c
~~~

There are seven pseudometabolites: lipid, protein, carbohydrate, DNA, RNA, cofactor, and ion.

To simulate the cell prioritising biosynthesis of lipids, we set the stoichiometric coefficients of all pseudometabolites except for lipids to zero in the above equation, giving:

~~~
     47.6 atp_c + 47.6 h2o_c + lipid_c
–> 47.6 adp_c + biomass_c + 47.6 h_c + 47.6 pi_c
~~~

Using this modified reaction as the objective function, the model was optimised using flux balance analysis, and this process was repeated for the other pseudometabolites, resulting in different growth rates for each round of ablation.

## Data availability

The data underlying this article are available in Edinburgh DataShare at https://doi.org/10.7488/ds/7845. The computer code underlying this article is available at https://github.com/arinwongprommoon/wongprommoonSinglecellMetabolicOscillations2024.

## Acknowledgements

AW acknowledges financial support from the Edinburgh Global Scholarship and the School of Biological Sciences, University of Edinburgh. AFM acknowledges support from the European Union’s Horizon 2020 research and innovation programme under the Marie Skłodowska Curie grant agreement no. 764591 (SynCrop). PSS acknowledges support from the BBSRC (grant number BB/W006545/1). We thank Ivan Clark for help, advice, and training; and Iseabail Farquhar for the CRISPR-Cas9 training required to construct insertion strains. For the purpose of Open Access, the authors have applied a CC BY public copyright licence to any Author Accepted Manuscript version arising from this submission.

