## Supplementary information for "Single-cell metabolic oscillations are pervasive and may alleviate a proteome constraint"

### Supplementary figures

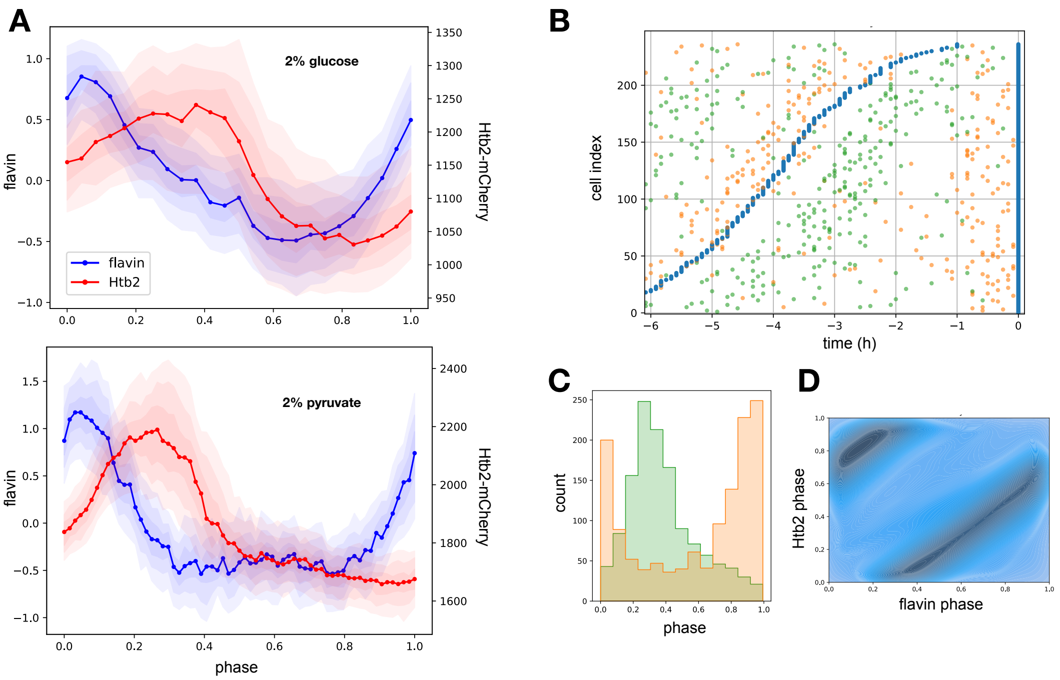


**Figure S1.** Flavin-redox oscillations in pyruvate are similar to those in glucose. **A** The waveforms of flavin (blue) and Htb2-mCherry (red) averaged over cells aligned at $t = 9$h, a time arbitrarily chosen, are similar in glucose and pyruvate. The flavin signal peaks at budding events and falls when Htb2-mCherry peaks, during M phase. Notice the longer G1 phase in pyruvate corresponds to a minimal flavin signal, approximately between phases 0.4 and 0.8. **B** A raster plot for one flavin oscillation shows that cell-cycle events occur in the same order relative to the flavin signal as they do in glucose; compare Fig. 1B. M phase, peaks in Htb2:mCherry, are in green; budding events are in orange. **C** Budding events and maximal Htb2 fluorescence occur at similar phases in the flavin cycle as they do in glucose; compare Fig. 1C. **D** A kernel-density plot of the flavin and cell-cycle phases shows phase-locked behaviour.

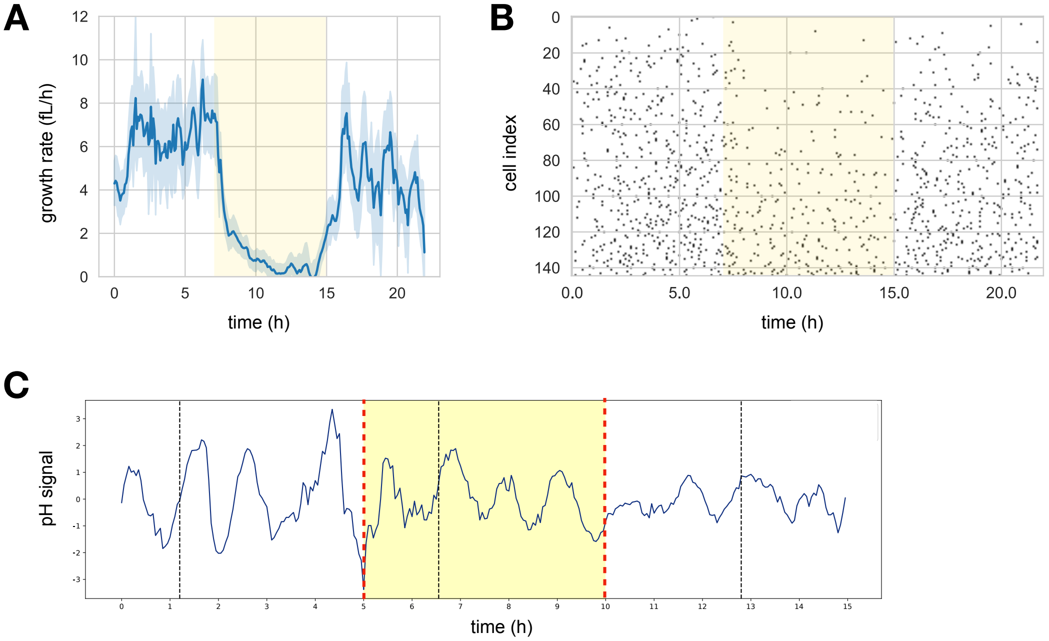


**Figure S2.** During starvation, cells slow their cell cycle but continue metabolic oscillations. **A** By switching from 0.75% glucose in minimal medium to just minimal medium from $t = 7$h to $t = 15$h, we slowed the cells' rate of growth. The mean growth rate, the mean rate of change of single-cell volumes, decreases almost to zero during the period of starvation (yellow shading). Data are from the experiment shown in Fig. 1E. **B** A heatmap of predicted budding events for the same cells shows that the density of budding falls during the period of starvation — compare with the period from $t = 0$h to $t = 7$h when cells are in 0.75% glucose. **C** An example time series from measuring the ratio of fluorescence emissions from pHluorin, which shows that intracellular pH continues to oscillate during glucose starvation. Here we switched BY4742 cells harbouring pHluorin from 2% glucose to 0% glucose in SC medium (yellow shading).

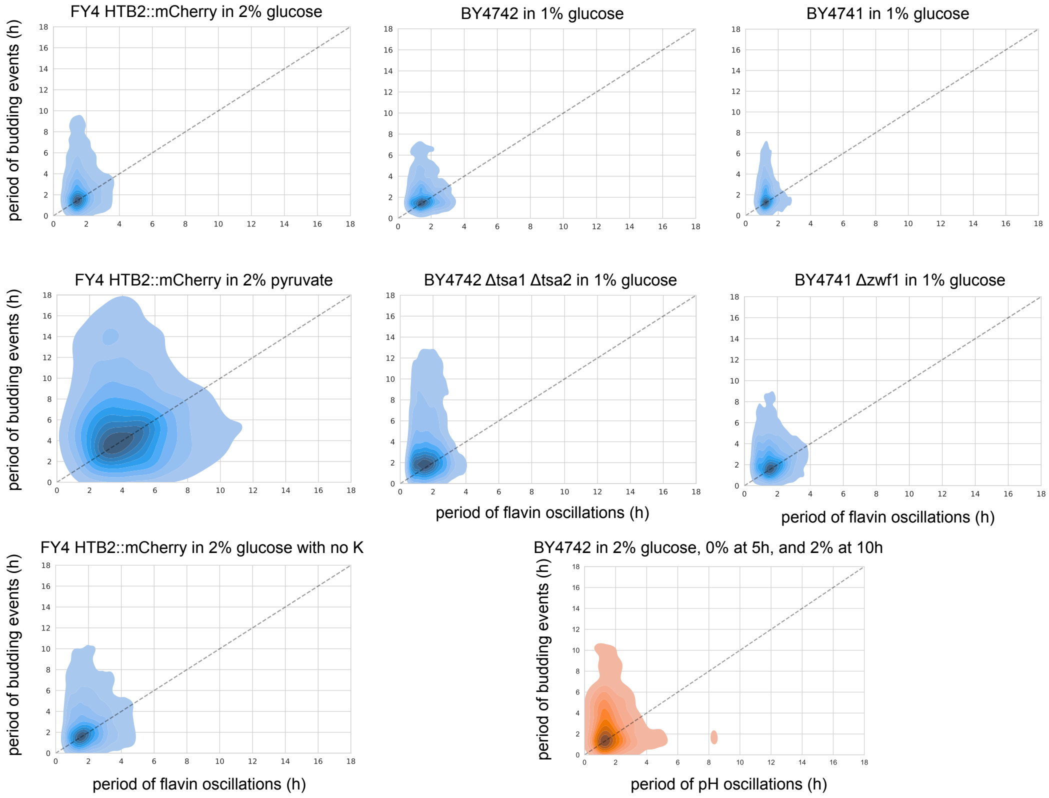


**Figure S3.** Metabolic oscillations have the same mean period as the cell cycle. We show kernel-density estimates found from plotting the two periods for all cell cycles and for all cells analysed in an experiment as points. We estimated periods by peak-to-peak times. Metabolic oscillations in flavin-redox state are in blue and in intracellular pH are in orange. The dashes show the $y = x$ line.

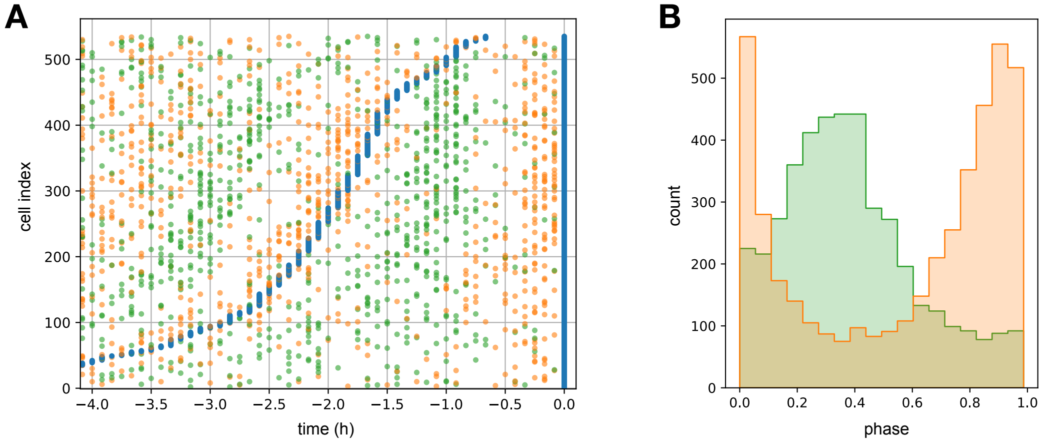


**Figure S4.** Eliminating potassium ions in the medium left the flavin-redox oscillations almost unaffected. **A** A raster plot for cells that have grown for eight hours in potassium-free medium. After six hours of growth, we switched cells into medium where we had replaced potassium phosphate with sodium phosphate. Data are taken from $t = 14$h. **B** Averaging over the entire experiment showed that the flavin-redox and cell cycles maintained approximately the same relative phase as they had in the presence of potassium; compare Fig. 1C. The experiment involved six hours of growth in normal minimal medium, followed by ten hours of growth in potassium free medium, followed by eight hours of growth again in minimal medium.

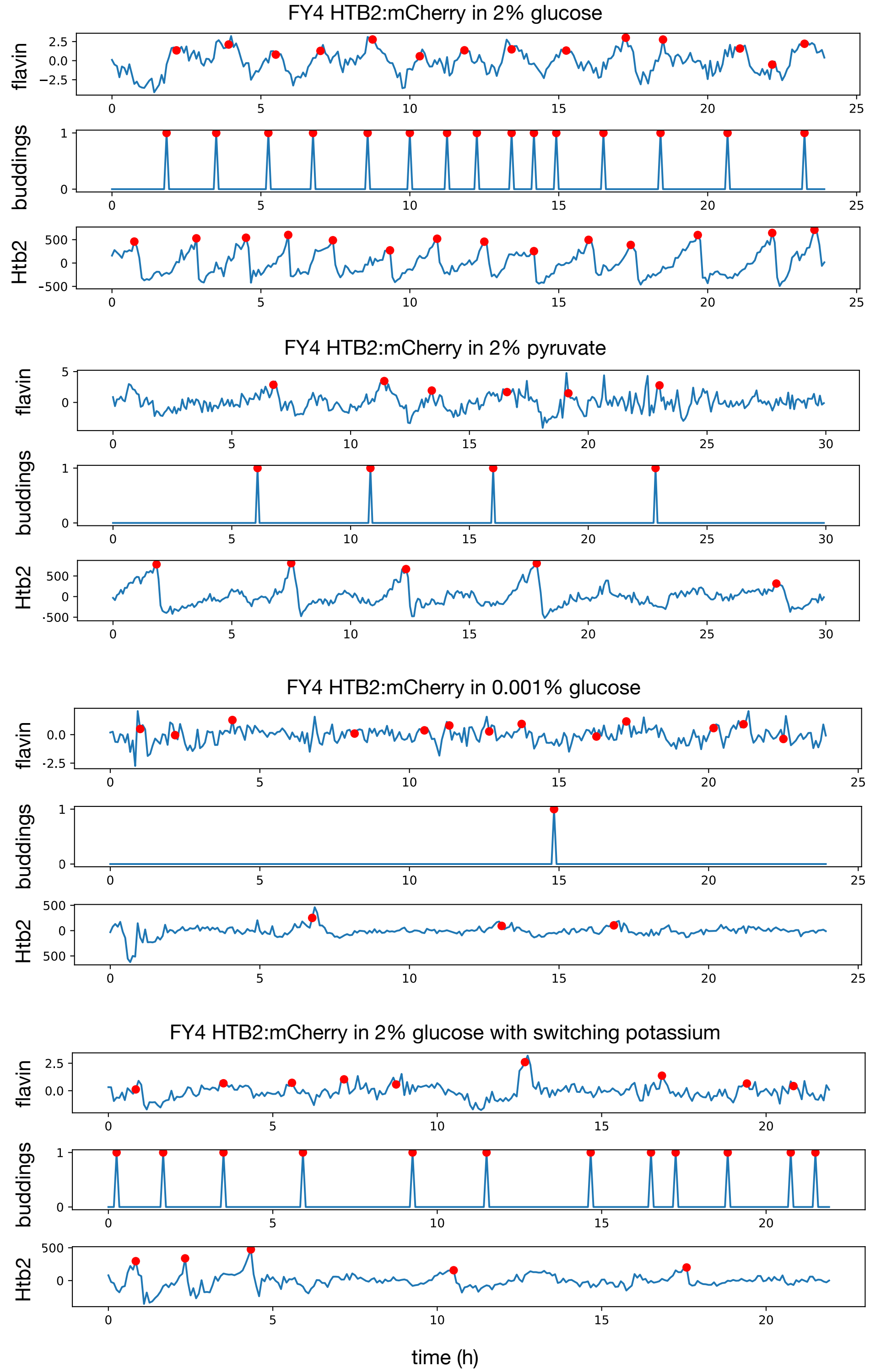


**Figure S5.** Examples of single-cell time series for the FY4 prototrophic strain in different media. Red dots mark automatically detected peaks in the signal, which we used to estimate periods. The low signal-to-noise rate in 0.001% glucose generated spurious peaks in the flavin signal. Budding events are predicted by the BABY algorithm (Pietsch et al., 2023).

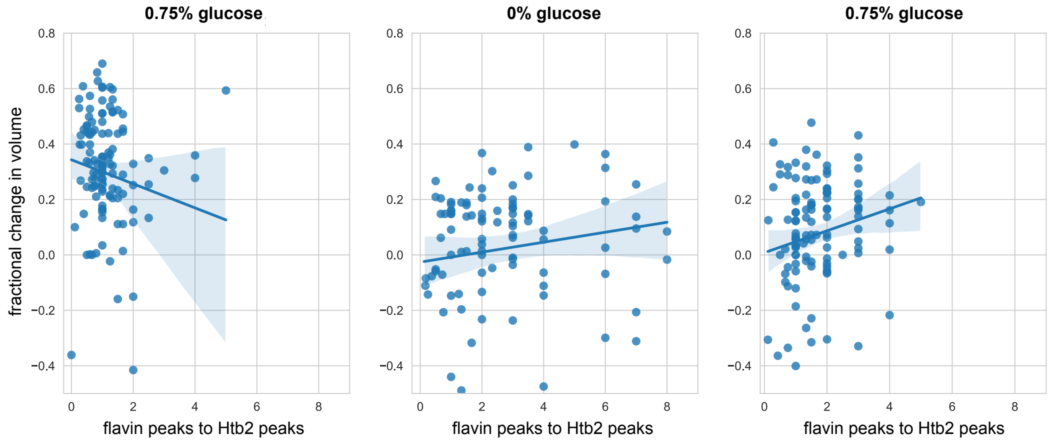


**Figure S6.** During starvation, there is weak evidence that cells with more flavin-redox oscillations per cell cycle have more growth. We divided the experiment in Fig. 1E into three stages: from 0h to 7h when cells were in 0.75% glucose; from 7h to 15h when cells were in 0% glucose; and from 15h to 21h when cells were again in 0.75% glucose. For each stage and for each cell, we found the change in volume over that stage, defined as the difference between the cell volume averaged over the last hour of the stage and the first hour of the stage, and plotted this difference against the ratio of the number of flavin-redox oscillations to the number of cell-cycle (Htb2) oscillations. For the first stage, in 0.75% glucose, most cells have equal numbers of flavin-redox and Htb2 oscillations, consistent with Fig. S3. There is a spurious negative correlation likely caused by outliers. For the next two stages — starvation and recovery from starvation — there is a positive relationship between the change in volume and the relative number of flavin-redox oscillations. The correlations though are weak: 0.14 for the second stage and 0.18 for the third stage.

### Supplementary methods

| Reagent | Concentration | Remarks |
|---|---|---|
| $KH_2PO_4$ | $3\,g\,L^{-1}$ | |
| $MgSO_4 \cdot 7\,H_2O$ | $0.5\,g\,L^{-1}$ | |
| $(NH_4)_2SO_4$ | $5\,g\,L^{-1}$ | |
| Trace metals | $1\,mL\,L^{-1}$ | See Table S2 |
| Vitamins | $1\,mL\,L^{-1}$ | See Table S3. Add upon use. |
| Carbon source | variable | Add upon use. |

**Table S1.** Composition of base minimal medium. For potassium-free media, replace $KH_2PO_4$ with $2.65\,g\,L^{-1}$ $NaH_2PO_4$, which gives the same molarity.

| Reagent | Formula | Concentration $[\mathrm{g\,L^{-1}}]$ |
|---|---|---|
| EDTA | $C_{10}H_{14}N_2Na_2O_8 \cdot 2\,H_2O$ | 15.00 |
| Zinc sulfate | $ZnSO_4 \cdot 7\,H_2O$ | 4.50 |
| Manganese (II) chloride | $MnCl_2 \cdot 2\,H_2O$ | 0.84 |
| Cobalt (II) chloride | $CoCl_2 \cdot 6\,H_2O$ | 0.30 |
| Copper (II) sulfate | $CuSO_4 \cdot 5\,H_2O$ | 0.30 |
| Sodium molybdate | $Na_2MoO_4 \cdot 2\,H_2O$ | 0.40 |
| Calcium chloride | $CaCl_2 \cdot 2\,H_2O$ | 4.50 |
| Iron (II) sulfate | $FeSO_4 \cdot 7\,H_2O$ | 3.00 |
| Boric acid | $H_3BO_3$ | 1.00 |
| Potassium iodide | KI | 0.10 |

**Table S2.** Composition of trace metal mix for minimal media described in Table S1.

| Reagent | Formula | Concentration $[\mathrm{g\,L^{-1}}]$ |
|---|---|---|
| D-(+)-biotin | $C_{10}H_{16}N_2O_3S$ | 0.05 |
| D-panthothenic acid calcium salt | $Ca(C_9H_{16}NO_5)_2$ | 1.00 |
| Nicotinic acid | $C_6H_5NO_2$ | 1.00 |
| *myo*-Inositol | $C_6H_{12}O_6$ | 25.00 |
| Thiamine chloride hydrochloride | $C_{12}H_{15}ClN_4OS \cdot HCl$ | 1.00 |
| Pyridoxal hydrochloride | $C_8H_{12}ClNO_3$ | 1.00 |
| 4-aminobenzoic acid | $C_7H_7NO_2$ | 0.20 |

**Table S3.** Composition of vitamin mix for minimal media described in Table S1.

#### Computing the mass fraction of each biomass component

##### Determining the molecular weights of pseudometabolites in ecYeast8

In ecYeast8, the objective function is defined via the biomass reaction:

```
47.5883 atp_c + 47.5883 h2o_c + lipid_c + protein_c + carbohydrate_c
+ dna_c + rna_c + cofactor + ion
-> 47.5883 adp_c + biomass_c + 47.5883 h_c + 47.5883 pi_c
```

with seven pseudometabolites: lipid, protein, carbohydrate, DNA, RNA, cofactor, and ion.

To obtain the mass fraction of each biomass component represented by the pseudometabolites, we treated each pseudometabolite as a chemical species and calculated its molecular weight by assuming mass balance (Chan et al., 2017; Dinh et al., 2022; Takhaveev et al., 2023). For the reaction producing the pseudometabolite — it is one of the products $p$, we therefore impose

$$\sum_{r=j}^{n_r} m_r c_r = \sum_{p=i}^{n_p} m_p c_p \tag{S1}$$

where $s = 1, \ldots n_s$ are the reaction's substrates; $p = 1, \ldots n_p$ are the products; $m_r$ and $m_p$ are the molar mass of reactant $r$ and product $p$; $c_r$ and $c_p$ are the stoichiometric coefficients of reactant $r$ and product $p$.

**Carbohydrate, DNA, RNA, cofactor, and ion pseudometabolites:** Computing the molecular weights of the carbohydrate, DNA, RNA, cofactor, and ion pseudometabolites is straightforward. Each of their equations has reactants with specified molecular weights and a sole product, the pseudometabolite. We can apply Eq. S1 directly, and the molecular weight of the pseudometabolite is $\sum_{r=j}^{n_r} m_r c_r$. The results are in Table S4.

| ID | Reaction | Computed molecular weight ($\mathrm{g\,mol^{-1}}$) |
|---|---|---|
| `r_4048` | `0.684535 (1->3)-beta-D-glucan + 0.228715 (1->6)-beta-D-glucan + 0.330522 glycogen + 0.650171 mannan + 0.126456 trehalose -> carbohydrate` | 350.37 |
| `r_4050` | `0.0036 dAMP + 0.0024 dCMP + 0.0024 dGMP + 0.0036 dTMP -> DNA` | 3.90 |
| `r_4049` | `0.0445348 AMP + 0.0432762 CMP + 0.0445348 GMP + 0.0579921 UMP -> RNA` | 64.04 |
| `r_4598` | `0.00019 coenzyme A + 1e-05 FAD + 0.00265 NAD + 0.00015 NADH + 0.00057 NADP(+) + 0.0027 NADPH + 0.00099 riboflavin + 1.2e-06 TDP + 6.34e-05 THF + 1e-06 heme a -> cofactor` | 4.83 |
| `r_4599` | `3.04e-05 iron(2+) + 0.00363 potassium + 0.00397 sodium + 0.02 sulphate + 0.00129 chloride + 0.00273 Mn(2+) + 0.000748 Zn(2+) + 0.000217 Ca(2+) + 0.00124254 Mg(2+) + 0.000659 Cu(2+) -> ion` | 2.48 |

**Table S4.** Straightforward cases of computing pseudometabolite molecular weights from pseudoreactions in ecYeast8.

**Protein pseudometabolite:** To compute the molecular weight of the protein pseudometabolite, we inspected reaction `r_4047`:

```
0.57284 Ala-tRNA(Ala) + 0.200644 Arg-tRNA(Arg) + 0.126979 Asn-tRNA(Asn)
+ ... + 0.330369 Val-tRNA(Val)
-> 0.57284 tRNA(Ala) + 0.200644 tRNA(Arg) + 0.126979 tRNA(Asn)
+ ... + 0.330369 tRNA(Val) + protein
```

In ecYeast8, aminoacyl-tRNA and tRNA species do not have specified molecular weights, and we treated `R`, which represents tRNA, as a chemical element of atomic mass 0 when computing $m_r$ for each reactant and $m_p$ for each product.

**Lipid pseudometabolite:** The lipid pseudoreaction is reaction `r_2108`:

```
lipid backbone + lipid chain -> lipid
```

but both `lipid backbone` and `lipid chain` have no specified molecular weight. Reaction `r_4065` specifies a lipid chain pseudoreaction:

```
0.0073947 C16:0 chain + 0.0217019 C16:1 chain + 0.0020726 C18:0 chain
+ 0.000796243 C18:1 chain -> lipid chain
```

which we use to calculate its molecular weight. Reaction `r_4063` specifies a lipid backbone pseudoreaction:

```
0.000631964 1-phosphatidyl-1D-myo-inositol backbone
+ 0.00243107 ergosterol + 0.000622407 ergosterol ester backbone
+ 0.000135359 fatty acid backbone + ...  -> lipid backbone
```

The model specifies the molecular weights in this reaction expect for `fatty acid backbone`. Four reactions produce `fatty acid backbone` (Table S5), and from each we estimated the molecular weight of `fatty acid backbone`. The values were slightly different, and we used their mean to calculate `lipid backbone`. We then found the molecular weight of `lipid` by summing the molecular weights of `lipid backbone` and `lipid chain`.

| ID | Reaction | Computed molecular weight ($g\,mol^{-1}$) |
|---|---|---|
| `r_3975` | `palmitate`<br>`-> 0.255421 fatty acid backbone`<br>`0.256429 C16:0 chain` | 742.54 |
| `r_3976` | `palmitoleate`<br>`-> 0.253405 fatty acid backbone`<br>`0.254413 C16:1 chain` | 744.56 |
| `r_3977` | `stearate`<br>`-> 0.283475 fatty acid backbone`<br>`0.284483 C18:0 chain` | 714.49 |
| `r_3978` | `oleate`<br>`-> 0.281459 fatty acid backbone`<br>`0.282467 C18:1 chain` | 716.51 |

**Table S5.** ecYeast8 reactions that generate the `fatty acid backbone` metabolite

We give a summary of the molecular weights in Table S6.

| Metabolite | Computed molecular weight ($g\,mol^{-1}$) | Biomass composition at growth rate $0.375\,h^{-1}$ ($g\,kg_{DW}^{-1}$) |
|---|---|---|
| Protein | 504.37 | 505 |
| Carbohydrate | 350.37 | 237 |
| RNA | 64.04 | 105 |
| Lipid | 31.57 | 57 |
| Cofactors | 4.83 | |
| DNA | 3.90 | 5 |
| Ions | 2.48 | |
| Total | 961.57 | |

**Table S6.** Computed molecular weights of bulk metabolites in ecYeast8, compared to an experimentally recorded biomass composition by Canelas et al. (2011).

##### Determining the mass fraction

We computed the mass fraction of each biomass component by dividing the molecular weight of the corresponding pseudometabolite by the molecular weight of biomass (Table S7).

| Metabolite | $f_i$ |
|---|---|
| Protein | 0.52 |
| Carbohydrate | 0.36 |
| RNA | 0.067 |
| Lipid | 0.033 |
| Cofactors | 0.0050 |
| DNA | 0.0041 |
| Ions | 0.0026 |

**Table S7.** $f_i$ values for each biomass component.
